# Image model embeddings for digital pathology and drug development via self-supervised learning

**DOI:** 10.1101/2021.09.20.461088

**Authors:** Khan Baykaner, Mona Xu, Lucas Bordeaux, Feng Gu, Balaji Selvaraj, Isabelle Gaffney, Laura Dillon, Jason Hipp

## Abstract

Whole slide images (WSIs) contain rich pathology information which can be used to diagnose cancer, characterize the tumour microenvironment (TME), assess patient prognosis, and provide insights into the likelihood of whether a patient may respond to a given treatment. However, since WSI availability is generally scarce during early stage clinical trials, the applicability of deep learning models to new and ongoing drug development in early stages is typically limited. WSIs available in public repositories, such as The Cancer Genome Atlas (TCGA), enable an unsupervised pretraining approach to help alleviate data scarcity. Pretrained models can also be utilised for a range of downstream applications such as automated annotation, quality control (QC), and similar image search.

In this work we present DIME (Drug-development Image Model Embeddings), a pipeline for training image patch embeddings for WSIs via self-supervised learning. We compare inpainting and contrastive learning approaches for embedding training in the DIME pipeline, and demonstrate state-of-the-art performance at image patch clustering. In addition, we show that the resultant embeddings allow for training effective downstream patch classifiers with relatively few WSIs, and apply this to an AstraZeneca-sponsored phase III clinical trial. We also highlight the importance of effective colour normalisation for implementing histopathology analysis pipelines, regardless of the core learning algorithm. Finally, we show via subjective exploration of embedding spaces that the DIME pipeline clusters interesting histopathological artefacts, suggesting a possible role for the method in QC pipelines. By clustering image patches according to underlying morphopathologic features, DIME supports subsequent qualitative exploration by pathologists and has the potential to inform and expediate biomarker discovery and drug development.

## 1 INTRODUCTION

In digital pathology, large quantities of time and money are spent annotating whole slide images (WSIs). This effort is the primary bottleneck for developing new deep learning tools and bringing these technologies into clinical practice.

*Self-supervised learning* [1] represents an opportunity to bypass this bottleneck. The idea, of which many flavours have been proposed in recent literature, is to train a model to predict parts of its input (e.g. words of a text or, here, pixels of an image) from other parts of its input. This approach allows to pretrain a model *without any annotation*, thereby benefiting from large amounts of unlabelled data. The resulting model is independent of any specific task: it does not know of any label, such as *cat* or *dog*. It builds, instead, a general-purpose internal representation of the structure of images: each image is internally converted into an *embedding*, that is a numerical vector that abstracts useful features of the image. Specific tasks such as classification can then be solved efficiently using the pretrained model as a feature extractor: given a set of images that are annotated for a specific task, e.g. classification, one can train a supervised model on the embeddings, rather than the initial images. The advantage is that the supervised model inherits from the pretrained model a representation of the images that is much richer than what could ever be learnt from the set of annotated images alone. This has proven to lead to significantly more robust models, especially when the annotations are expensive and scarce [2–5].

In the context of histopathology, embeddings obtained through self-supervised learning have the potential to solve a broad number of problems. They can serve as model feature inputs to be used for downstream supervised learning tasks such as image classification and survival prediction. Embedding images into an *N* -dimensional space additionally has the advantage that the distance between points in this space can provide useful information on the semantic similarity between the corresponding images; as we will discuss, this opens the way for a range of tools that can either offer new insights based on the analysis of clusters, or enable novel functionality as diverse as automated patch annotation, image similarity search, or patch quality control.

This potential is, however, largely untapped, and the application of self-supervised methods in histopathology is at a very preliminary stage compared to the work on “natural” images (See Section 2). Applying deep learning to WSIs, specifically Haematoxylin & Eosin (H&E) stained WSIs, raises a number of challenges:

1. Directly applying standard image processing deep learning approaches to WSIs is difficult because WSIs are generally too large to be processed directly by a neural net.
2. Pretrained models are most commonly trained on Imagenet [6], and other databases of ‘natural’ images, which are not only of very different size compared with histopathology images, but also include very different types of objects.
3. WSIs encode image data at multiple zooms & resolutions, thus either some choices must be made about the level of interest, or a strategy must be implemented to encode multiple levels effectively.
4. WSIs are commonly labelled at the slide level, and not at the patch or pixel level (e.g. with patient outcome), so embeddings generated at the patch level must later be aggregated if used for slide level predictions.
5. WSIs collected from different medical centers typically exhibit differences in terms of scanner characteristics and staining protocols. For example H&E staining is routinely used to differentiate nuclear from cytoplasmic cellular components; however the depth of the purple hue it imparts is generally of little interest.

In the field of drug development these challenges are even more severe. There are few publications demonstrating deep learning models applied to clinical trial datasets curated for novel drug development, and perhaps none utilising unsupervised or self-supervised learning approaches. Almost certainly this is due to the extreme scarcity of data as clinical trials, especially early stage ones, may enroll just 10s of patients, far fewer than is generally required to train and test deep learning models. Overcoming these challenges to enable the extraction of information from tissue imaging, even from small trials, is crucial for advancing drug development to bring new therapies to patients.

Beyond the aforementioned applications of image embeddings to histopathology (i.e. survival prediction, patch quality control etc.), in drug-development there is the further opportunity to perform exploratory analysis of the embedding space, identifying common morphologic features across non-responsive patients. Such exploratory work allows a pathologist to quickly discover dataset-wide or even cross-dataset patterns, providing new opportunities for hypothesis generation that can steer future drug development opportunities.

In this work we describe the construction and training of a deep learning pipeline for generating self-supervised embeddings of image patches from H&E-stained WSIs called DIME (Drug-development Image Model Embeddings), including our approach to addressing each of these technical challenges. Additionally, we compare two self-supervised approaches - inpainting and contrastive learning - for clustering (on TCGA and Camelyon16) and classifying tumour patches (on Camelyon16 and Mystic, an AstraZeneca-sponsored Phase III clinical trial), and qualitatively explore the resulting clusters to derive insights on pipeline performance. Notably, we demonstrate the application of DIME to Mystic, which is a multi-site, global study with substantial variability in tissue and staining.

## 2 RELATED WORK

As digitized H&E WSIs are becoming increasingly abundant, many endeavours have been made to train deep learning models using these data. For a general overview of existing work, [7] gives a survey of relevant approaches.

### Metric Learning and Similarity Search

The DIME pipeline is inspired by the work on SMILY [8], where a substantial effort was made to produce embeddings capturing image similarity for H&E WSIs. SMILY is based on a convolutional neural network trained on about 500,000,000 “natural images” (e.g. dogs, cats, trees, man-made objects) from 18,000 distinct classes. The pretrained model is used to map input WSI patches into the embedding space for carrying out similarity search. SMILY utilises a *metric learning* algorithm, called trinet loss [9, 10], to produce embeddings. This approach works by pairing images into triplets of anchor, positive, and negative; in effect domain knowledge is used to provide labels that indicate which images are more ‘similar’, according to the domain expert. It assumes a clear understanding of what constitutes ‘similarity’ by the domain expert, and hence biases the embeddings towards certain types of similarity, and away from others.

The authors of [8] used this approach to develop a powerful image similarity model for H&E WSI patches; in this case ‘similarity’ was constituted by images being of the same tissue type or having similar histopathological features; but this required a substantial annotation effort by pathologists to make this image pairing possible (127,000 patches from 45 WSIs). The data were from TCGA, which contains more than 11,700 WSIs; so it is clear that even with this enormous labelling effort, several orders of magnitude more data are available for training, though unlabeled. Furthermore, since metric learning algorithms require similarity to be effectively defined by human users’ annotations, the resultant embeddings are likely to be less suitable for future downstream predictive models (e.g. for survival prediction of some new therapy).

For the DIME pipeline we wanted to take a more generic approach, producing embeddings that would be useful for a wide range of downstream applications from image similarity search, to automated annotation, quality control, and for training down-stream classifiers. Here another class of algorithms, known as *self-supervised* learning, is a more appropriate choice, as it does not rely either on labels or image pairings. In self-supervised approaches a meta-problem is posed around the input data itself that is expected to yield embeddings which can broadly understand the structure of the data. In doing so, ‘similarity’ is not defined by domain expert labelling, but by the characteristics of the objective function of the learning algorithm. It is therefore reasonable to expect such algorithms may be able to extract previously unidentified features in the images which are more broadly applicable to a variety of down-stream applications. Since they also do not require annotations for image pairing, it is possible to train on a larger set of TCGA data with the primary constraint being the cost of computation, and not the cost of expert pathologists’ time.

### Self-Supervised Learning Techniques

One example of a self-supervised learning algorithm is image inpainting [3], where a model learns to reconstruct images from which parts have been blanked out. Such models have demonstrated a powerful ‘imagination’ for filling in missing data, restoring corrupted images, or achieving super-resolution [11]. Succeeding at such tasks implies that the trained embeddings must be capable of understanding, from surrounding context, what constitutes a plausible looking image; it is not such a stretch, then, to assume that such embeddings may be generically useful for a range of downstream tasks (image similarity, automated annotation, prediction, etc.).

Another approach to self-supervised learning is contrastive learning that explicitly aims to learn representations (i.e., an embedding space) in which similar images stay close to each other while dissimilar ones are far apart. One popular implementation of this is SimCLR [3], which has shown good performance at image classification compared with supervised approaches. In SimCLR, images and their augmented versions constitute positive pairs (and hence should appear near each other in embedding space), whereas different images constitute negative pairs (and hence should appear relatively further from one another in the embedding space).

### Self-Supervised Methods for Histopathology

Several previous attempts have been made to train self-supervised learning models on histopathology images, with varying degrees of success. Most commonly these attempts have utilised the Camelyon16 dataset, along with one or two others, giving the learned model exposure to only a very small range of cancer types, as in [12–15].

[16] stands out as an exception to these more narrowly trained models, whereby an impressive 60 histopathology datasets were utilised for model pretraining. The authors report a surprising finding: contrary to the natural image domain, training on a tiny fraction (less than 1%) of the dataset did not negatively impact the resulting model. The authors explain this outcome by noting that training on an increasingly large quantity of data tends to decrease the relative diversity of the data per patch. While we appreciate the crucial role of data diversity, we expect that with careful attention to promoting diversity, greater performance can be obtained by dramatically increasing the quantity of data used.

[13] also stands out amongst the prior work. Although the authors trained on datasets covering a relatively narrow set of cancer types, the SimCLR approach combined with multiple instance learning seems to be a very powerful approach for pretraining embeddings for WSI classification models. This paper focuses on boosting the performance of whole slide image classification by using the same dataset for SimCLR pretraining as used for the downstream classification task. We note, however, that for practical application to drug development the downstream dataset will be far too scarce to train a self-supervised model. This raises the need for pretraining one self-supervised model that is general enough to abstract features for a wide range of histopathology images, with relatively limited downstream training required.

In [17] the desire to produce general histopathology embeddings was highlighted, and implementations of contrastive predictive coding were trained to attempt this. The authors concluded that this high bar was not achievable with this type of learning algorithm and model since they observed that only the quickly learned low-level features were utilised for downstream tumour classification. While this result is disappointing, it does not demonstrate that other self-supervised learning algorithms will not succeed where contrastive predictive coding failed.

## 3 CONTRIBUTIONS

In this paper we document contributions towards an image model embedding pipeline for drug development that seeks to capture features that are generally applicable to a wide range of downstream applications, and can be directly applied to real world clinical trial datasets obtained in drug development studies. The key differences between this work and the prior literature are: First, we utilise a training dataset from TCGA containing orders of magnitude more data, as well as more diverse data. Second, we train inpainting and SimCLR algorithms; SimCLR is known to compare favourably with many of the self-supervised algorithms evaluated in prior work, while inpainting has not, to our knowledge, been applied to the histopathology image embeddings domain. Thirdly, we explore how the pretrained embeddings can be utilised for drug development focused applications, and address the particular technical challenges this imposes.

Specifically we document the following contributions:

- We propose a novel deep-learning based pipeline called DIME (Drug-development Image Model Embeddings) for extracting generic, multi-scale image features that enable similarity search, quality control and patch classification for whole slide images, effective even for small datasets.
- We demonstrate that a model trained on a large and diverse dataset can be used to produce relevant embeddings for downstream applications on unseen datasets. In particular we applied the model trained on TCGA, to Camelyon16 and a dataset from an AstraZeneca-sponsored Phase III clinical trial.
- We identify the relative strengths of inpainting and SimCLR. Specifically inpainting outperforms SimCLR at capturing histopathologic features aiding in tissue/disease-type clustering. Conversely, SimCLR is more robust to colour variations introduced by staining and scanning.
- We outline the application of DIME to digital pathology for drug development scenarios including the improvement of pathologist workflow, functioning as a QC tool prior to the application of other analysis methodologies, and classification tasks to relate patho-morphologic information contained in WSIs with clinical or translational measurements (e.g. outcome, mutation status, gene expression).

## 4 DIME PIPELINE

In this section we describe the initial construction of the DIME pipeline, detail its training on a large quantity of TCGA data for building a generic feature extractor which inferences embeddings, and outline the process for evaluating embeddings in downstream applications.

### 4.1 DIME Pipeline construction

Here we describe the pipeline from input WSIs to trained image patch embeddings, and suggest optional downstream pipeline stages for making use of these embeddings (e.g. for patch similarity search as in [8], or for downstream patch classification), as shown in Figure 1.

**Figure 1:**
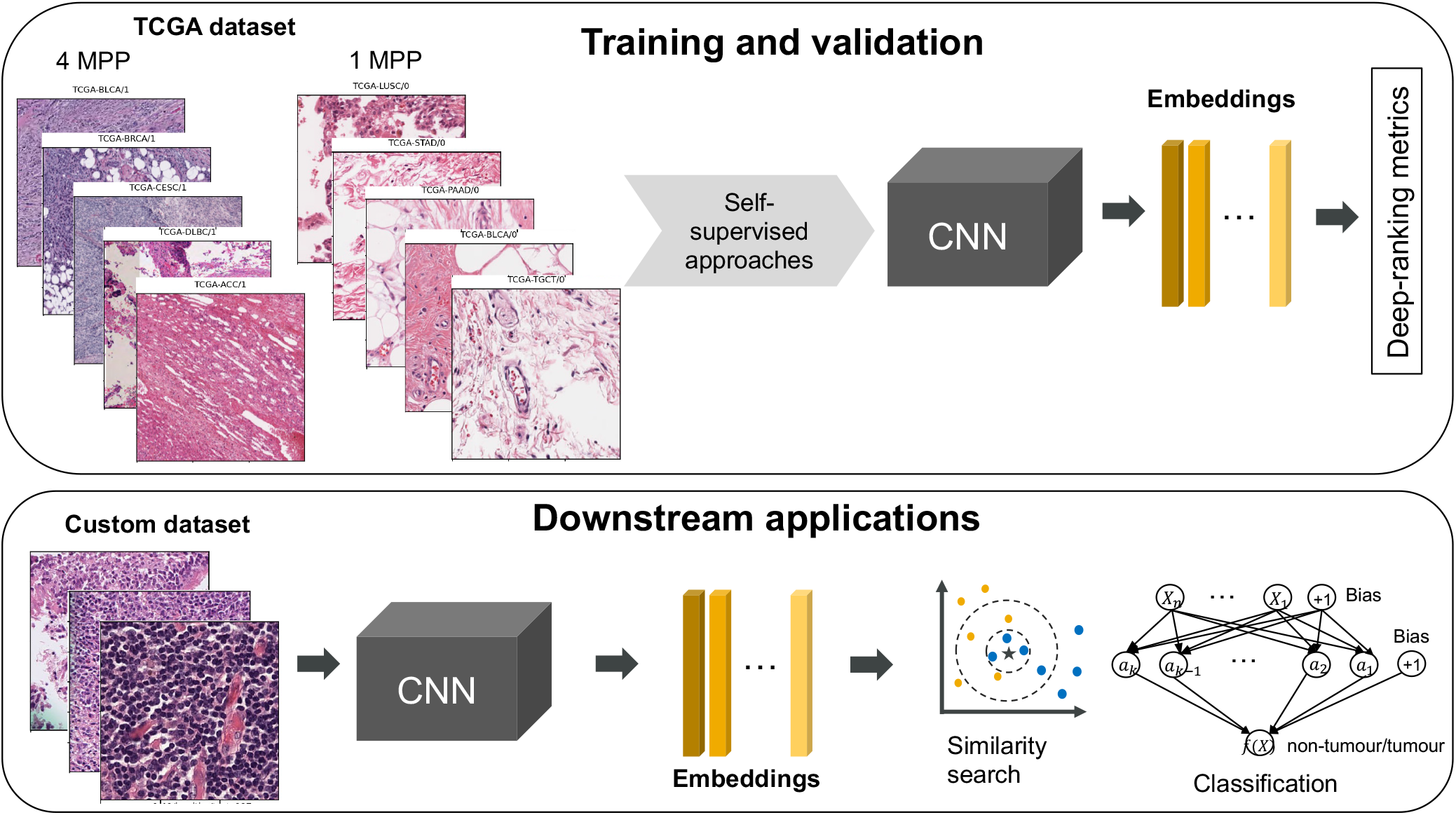
Diagram of DIME pipeline.

#### 4.1.1 Data ingestion

We downloaded 2,962 diagnostic WSIs from National Cancer Institute Data Portal ^1^, stratified across 32 projects (investigations). These projects provide slides characterising 32 cancer types, including 10 rare cancers. The selection process was to utilise 100 images per project (for each project with at least 100 diagnostic slides, and all available diagnostic slides for projects with fewer with the fewest being 39). This approach ensured reasonable diversity of projects (e.g., cancer types), which we expect to benefit the generalisability of the final trained feature extractor.

We split the dataset into 2836 slides for training and validation, and 126 slides for testing. The testing slides cover 29 TCGA projects. The 2836 slides reserved for training and validation have disjoint slide IDs.

We also utilised a reduced, but diverse, subset of the TCGA data. The diverse TCGA dataset is a subset of the larger dataset of 2962 slides and contains 94 slides for training and 20 slides for validation. The diversity of the TCGA slides is characterised by the combination of primary site (e.g. breast) and disease type (e.g. adenomas and adenocarcinomas). There are in total 94 combinations (classes) and only one slide is selected from every class to form the training subset of the diverse TCGA dataset. Similarly, the 20 validation slides come from 20 classes that have the most cases. We use the same 126-slide testing set in all cases which has 37 combinations of primary site and disease type.

#### 4.1.2 Tissue filtering and patching

In order to train on the WSIs, it is necessary to first extract image patches. In part this is because WSIs are generally too large at the highest resolution to fit as an input for standard image processing deep learning models such as ResNet34; but also because WSI formats generally store the image at multiple levels of zoom & resolution to assist pathologists with exploration at multiple levels of detail. Based on the recommendation of pathologists involved, 1 and 4 microns per pixel (MPP) were considered levels of zoom at which pathologists could make meaningful interpretations of tissue structures. MPP levels vary across diagnostic slides within the TCGA however, so we used a heuristic of selecting the two closest MPP levels per image to 1 and 4 MPP. Naturally, this results in a non-uniform distribution of MPPs across training data; however we believe that the more populous zoom and resolution levels broadly represent appropriate levels for pathologic inspection, and the non-uniformity across images assists model generalisability by enhancing the diversity of data that the embeddings must train to account for, see Figure 10. Patches at a fixed size of 224 by 224 pixels, selected to match the input to ResNet34 - which we would be using as part of our model architecture.

From each of the digital WSIs, we extracted patches of 224 × 224 pixels from the two levels closest to 1 MPP and 4 MPP, corresponding to roughly 10x and 2.5x magnification levels, respectively. Hence, every patch is linked to a particular slide ID and MPP level. Note that the resulting patches from one slide could contain some tumour or all non-tumour type of tissues.

The training subset yields 7,691,128 patches at 1 MPP and 535,074 patches at 4 MPP. The validation subset yields 428,645 patches at 1 MPP and 38,875 patches at 4 MPP. The test subset yields 374,989 patches at 1 MPP and 26,318 patches at 4 MPP.

Tissue filtering was also applied in order to minimise computation time wasted training on entirely blank background slides. Given the large quantity of data to be processed, we opted for a relatively simple but efficient tissue filtering algorithm known as OTSU [18]. We chose a tissue proportion cutoff of 10%, excluding any patches with less than this fraction of tissue contained within.

Since tissue sizes are not constant across diagnostic slides, WSIs each resulted in a different quantity of patches ranging from 28 to 7487 at 1 MPP and from 3 to 1557 at 4 MPP in the selected TCGA dataset described in Section 4.1.1. In order to mitigate this vast range of patches per WSI, we use an upsampling procedure whereby an equal number of patches per slide (and per MPP) are used for training.

Given the sheer scale of the data (total 8,693,722 patches across 2,836 WSIs) it is extremely difficult to apply any manual quality control to the input data. One deleterious effect that was identified is that images with pen-marking, produced when pathologists draw on glass sides with a pen as part of their annotation process, can sometimes be incorrectly filtered by the OTSU algorithm, resulting in fewer than expected patches from some images. While we look to improve upon this in future work, we do not consider quality controlling the input data to be of great importance here; this is because when the DIME pipeline is working correctly we expect image patches of poor quality or with artifacts such as pen markings to be clustered in embedding space. This is confirmed by our observations of the embeddings produced for several datasets we experimented with, see e.g. Fig. 9. In other words, part of the value of the DIME pipeline is to offer a mechanism for quickly quality controlling images since any distinct similar image corruption should cluster in embedding space. On this basis, one could make the argument for not including tissue cutoff filtering at all, and simply including all patches regardless. While this approach should work, it incurs a substantial additional cost in processing a vast majority of completely blank “background” patches from slides which we largely avoid here.

#### 4.1.3 Self-supervised learning algorithms

We initially trained a feature extractor using the self-supervised inpainting approach on the TCGA dataset described in Section 4.1.1, but after careful examination of the patches in our embedding space, especially of embeddings produced on the Camelyon16 dataset (see section 6), we became concerned that our trained inpainting model may be too sensitive to these colour based factors. A significant problem in training embeddings for querying similar histopathology images is whether irrelevant characteristics of the images may dominate the embedding space. For H&E-stained WSIs, a serious concern is that the particular staining or scanning of an image may dominate, despite being of generally little interest compared with morphologic features. Contrastive learning approaches are expected to be more robust to variations in staining and scanning, because by specifying the augmentations used during training the embeddings can explicitly ignore certain variations across the data. We therefore additionally trained a SimCLR model as an alternative against which to compare.

##### Inpainting

For the inpainting model architecture we adopted a U-Net [19] based on a ResNet34 encoder. This allows us to perform transfer learning by utilising a ResNet34 model with weights pretrained on ImageNet data [6]. The decode arm of the U-Net is also, therefore equivalent to the reverse ResNet34 architecture but including relevant concatenation stages at each level. This architecture is similar to the one presented in [20] and our implementation, similarly, uses the FastAI library [21].

The inpainting approach comprises of filling blacked-out boxes overlaid on the input images, with details in Section 8. Intuitively, this task should result in embeddings that broadly capture information about the structure of the WSI patches (since the task is to fill in missing data).

Rather than rely directly on pixel errors, we take an approach suggested in [22] and append a VGG16 model (pretrained on ImageNet) to the output of our U-Net. The objective function is then defined by the combined pixel loss, feature loss, and style loss. Each of these has a different role to play in arriving at perceptually similar patches. Naturally, the pixel distance is relevant, but often attributes equal weighting to types of error that are perceptually extremely unequal. Feature and style losses capture the broader similarities between higher level structures (effectively captured at increasing levels in the pretrained ImageNet model). While the pretrained VGG16 ImageNet model has been trained on large quantities of naturalistic images, rather than on histopathology images, the intermediate layers nevertheless tend to detect simplistic shapes (such as lines, circles, and diagonals) that are the building blocks of both naturalistic images and histopathologic structures - thus using feature and style loss via such a model still seems to be effective.

Given an input image with black boxes *I*_*in*_, the network prediction *I*_*out*_, and the ground truth image *I*_*gt*_, we first define our per-pixel losses as the L1 losses on the network output:

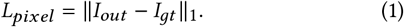

Then, we define the perceptual loss similar to that in [23], as the L1 distances between *I*_*out*_ and the ground truth *I*_*gt*_, but after projecting these images into higher level feature spaces using a VGG-16 model pretrained on ImageNet:

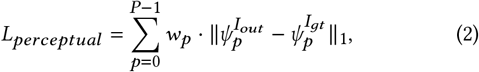

where 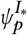 is the activation map of the *p*_th_ selected layer given original input *I*_∗_. We use layers 22, 32 and 42 for our loss. Through a small scale hyper-parameter exploration, we identified 5, 15, 6 to be good choices for *w*_*p*_ to weigh the loss from the three selected layers.

The third component of the loss function is the style-loss term, which is similar to the perceptual loss but uses an autocorrelation (Gram matrix) on each feature map before applying the L1 loss:

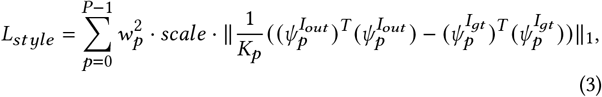

where the same weight *w*_*p*_ is applied to the *p*th selected layer in the style loss as in the perceptual loss. We choose *scale* of 5*e*3 for scaling the style loss versus the perceptual loss. Given the high-level features 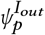 of shape *C*_*p*_ ∗ *H*_*p*_ ∗ *W*_*p*_, we apply the normalization factor 1 /*K*_*p*_ = 1 /*C*_*p*_*H*_*p*_*W*_*p*_ for the *p*th selected layer.

The total loss is a combination of the pixel loss, perceptual loss, and style loss,

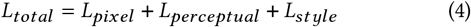

##### SimCLR

Our SimCLR architecture design follows that introduced in [3] which uses an encoder followed by a MLP with one hidden layer (and ReLU activation) as the projection head that maps the representations to the latent space. We chose ResNet34 as the encoder architecture that matches with the encode arm of our in-painting models, but with two subsequent fully-connected layers, each with 512 neurons.

Under SimCLR, the pipeline applies random data augmentation to generate two correlated versions of each data sample, which are considered a positive pair. For each positive pair the other augmented samples in the minibatch are treated as negative samples. The contrastive loss uses the normalized temperature-scaled cross entropy loss as introduced in [3].

After training, the projection layers are removed since the experiments carried out in [3] show the layer before the projection head gives a better representation than the projection layers. Therefore, the embeddings captured from the encoder are used for downstream analysis. By using the embeddings from the same encoder layer, it gives a fairer comparison between the architectures used for in-painting and SimCLR. To train SimCLR, we initialise with weights from the ImageNet-ResNet34 model. More details of training setup can be found in Section 8

#### 4.1.4 Model variants

Given the inpainting and simCLR algorithms, we present here the five variations of the DIME pipeline trained on a large or reduced TCGA dataset for comparison:

(1) DIME-Inpainting
(2) DIME-Inpainting-D
(3) DIME-Inpainting-DCT
(4) DIME-SimCLR-DC
(5) DIME-SimCLR-DCA

In later evaluations we additionally include:

(6) ImageNet-ResNet34: ResNet34 pretrained on imagenet
(7) SimCLR-D: a SimCLR model^2^ pretrained on 400,000 diverse medical images (hence D for diverse) that are sampled from 26 non-WSI datasets and 35 WSI datasets, as described in [16],

for benchmarking purposes.

##### DIME-Inpainting

The DIME-Inpainting model was trained using the set of 2836 training and validation slides from TCGA. Notably, the colour augmentation applied to the image patches when training this model is very light with a hue factor of 0.02, a saturation factor of 0.1, a contrast factor of 0.2 and a brightness factor chosen uniformly from (0.4, 0.6).

##### DIME-Inpainting-D

The DIME-Inpainting-D model differs from the DIME-Inpainting model only in that it was trained using the reduced size diverse dataset of 94 training slides (243,083 patches) and 20 validation slides (72,072 patches), hence D for diverse. Test performance is reported on the identical set of 126 slides in all model variants. More details are given on the diverse dataset in 5.1).

##### DIME-Inpainting-DCT

The DIME-Inpainting-DCT (DCT for diverse, colour augmented, target adjusted) model was also trained on the diverse dataset (hence D for diverse), and further adapted in two ways. Firstly, a stronger colour augmentation is utilised with a hue factor 0.06 and a saturation factor of 0.25 (hence C for colour augmented). Secondly, the target image which the UNET aims to generate is in this case set as the image prior to hue and saturation augmentations, but after morphological augmentations (hence T for target adjusted).

##### DIME-SimCLR-DC

The fourth model variant is based on the simCLR approach, denoted as DIME-SimCLR-DC, which utilised the diverse dataset (hence D for diverse) with colour augmentation the same as DIME-Inpainting-DCT (hence C for colour augmented).

The DIME-SimCLR-DC model uses all the same data augmentations as in the DIME-Inpainting-DCT case, also utilizing the a hue factor 0.06 and a saturation factor of 0.25.

##### DIME-SimCLR-DCA

The fifth and final model variant, DIME-SimCLR-DCA, utilised the diverse dataset with very aggressive colour normalisation (hence A for aggressive colour augmentations), which matches that used in [3] with a hue factor of 0.2, a brightness factor of 0.8, a contrast factor of 0.8 and a saturation factor of 0.8.

### 4.2 Model Evaluation

When attempting to train embeddings that are generically applicable to a wide range of downstream tasks, identifying an appropriate measure of performance is a non-trivial consideration. For example, one might identify performance across multiple downstream tasks, but then it may be unclear as to how (or whether) weighting should be applied across these tasks, and this furthermore does not account for future downstream tasks that may later present themselves. It is preferable, perhaps, to identify some positive intrinsic characteristics of the trained embeddings that provide evidence of their value.

If the WSIs have labels with which they may be clustered, the effective clustering of the patches according to those labels may be used as an indication of performance. Any such labels would have to be a proxy for some notion of ‘similarity’, as previously discussed with reference to metric learning algorithms, and any such metric should therefore be considered as indicative of performance, rather than definitive. The metric learning literature offers a useful insight into an appropriate evaluation metric for the performance of embeddings. In [24] the weaknesses are highlighted for some common evaluation metrics such as Recall@K, Normalized Mutual Information (NMI), and the F1 score. Mean Average Precision at R (MAP@R) is proposed as a combination of the ideas of Mean Average Precision and R-precision. MAP@R is designed to reward well-clustered embedding spaces and proven to be more informative and stable than Recall@1.

For TCGA we opted to use the metadata for primary site and disease type together with MPP level as labels against which to compute the MAP@R. Labelling by combined tissue and disorder type, as well as by zoom, captures most of the distinct classes for which we expect clustering. However, from the same slide there can be tumour and non-tumour patches, which have distinct histopathological features; such cases result in reduced evaluation score despite good clustering performance by the algorithm.

This imperfection of the MAP@R metric is worth dwelling on, since there are additional reasons to be distrustful of any such objective clustering metric. As already noted, tissue type is not the only valid and relevant sense in which images may be ‘similar’. In our experimentation we noticed embedding clusters that included multiple overlapping TCGA cancer types, but which, upon examining the patches visually, had good reason to be clustered; for example, image patches may have:

- Image corruptions or artefacts, such as blurring (implying that the embedding training pipeline is useful for quality control).
- A substantial quantity of slide background even though the slide passed the 10% tissue filtering cutoff.
- Histopathology artefacts which may occur in multiple tissue types.

Conversely, we also identified multiple separate embedding clusters for data from the same project. This was most commonly due the presence of image patches at multiple different magnifications.

Despite these limitations, we found that MAP@R still offered a good approximate indication of the clustering of embeddings, which is a meaningful objective proximate measure of embedding performance. It is necessary, however, to supplement MAP@R with a subjective exploration of the embedding space in order to understand what types of errors are occurring. Additionally, we trained downstream classifiers using these embeddings to demonstrate the capability of the DIME pipeline for capturing relevant morphopathologic features (see section 6).

## 5 PRETRAINING ON TCGA

In this section we explore the DIME pipeline model variant embeddings and report objective evaluation metrics for the TCGA dataset.

### 5.1 Pretraining approach

We trained every DIME model in a distributed setup across 4 GPUs. We derive the 512-dimensional embeddings from the encoder on the validation subset and then calculate the deep-ranking metric, MAP@R. Rather than training on full epochs, models were trained on TCGA for a maximum of 200 *rounds*, with each round comprising 100,000 patches. We used early stopping (with details in Section 8) based on weighted average MAP@R, with a patience of 20 rounds. DIME-Inpainting model training was the exception, which due to the much larger quantity of data was trained for 1600 rounds.

We calculated both the average and the weighted average MAP@R metrics, where the per-sample MAP@R is defined in [24]. The weighted average MAP@R gives the average of MAP@R across all samples regardless of their class, while the average of the average MAP@R gives the average of MAP@R per class. When there is class imbalance, the weighted average MAP@R will be dominated by the classes that contain more samples. The metric has a parameter *R* whose value is determined by the number of samples in the smallest class, which is 17 from the class ‘Esophagus/Adenomas and Adenocarcinomas/4MPP’ in this case. It is noted that TCGA slides do not have exhaustive tumour annotations, so the metric reported here does not accurately reflect the similarity between tumour or non-tumour patches.

### 5.2 Validation of the TCGA Training

Here we explore how well the pretrained TCGA embeddings cluster, and compare the DIME model variants against the ResNet34 pretrained on ImageNet, which is referred to as ImageNet-ResNet34 and used as a baseline model.

#### 5.2.1 DIME-Inpainting TCGA UMAP

Uniform Manifold Approximation and Projection (UMAP) [25] down-projections of embeddings for the TCGA test images are shown in Figure 2 for the DIME-Inpainting-DCT model alongside the ImageNet-ResNet34 model. Here the embeddings are labelled according to TCGA project and MPP level, with each project indicating a different cancer type (over many different tissue types).

**Figure 2:**
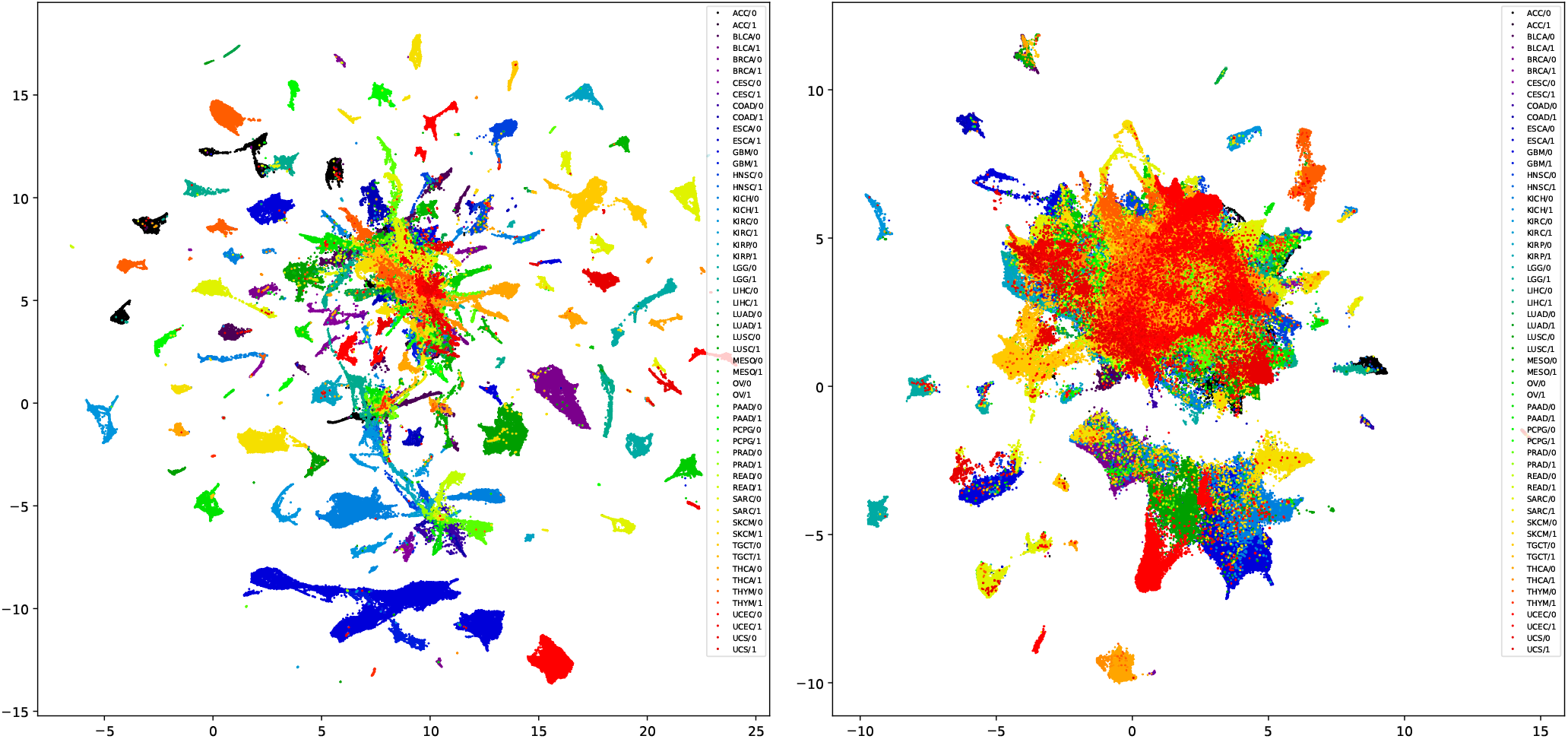
UMAP visualization of TCGA test images for DIME-Inpainting-DCT (left) and ImageNet-ResNet34 (right). The DIME embeddings show dramatically superior clustering on tissue and disease type over the imagenet model.

For the DIME-Inpainting-DCT model the down-projected embeddings show good clustering according to cancer type and MPP level, with many separate and distinct clusters visible. Subjectively, it appears that the embeddings from the DIME-Inpainting-DCT model have superior class discrimination compared with ImageNet-ResNet34 model for the TCGA dataset. UMAPs for other DIME variants performed similarly, with the contrastive approach showing slightly poorer clustering.

#### 5.2.2 Results of Model Evaluation

Here we report MAP@R as an objective metric of clustering according to tissue, disease type, and MPP level. The metrics, shown in table 1, support the subjective impression that the DIME-Inpainting models produce the best separation by tissue and disease type. The three DIME-Inpainting variants have the highest MAP@R scores, with DIME-SimCLR-DC not too far behind. The ImageNet-ResNet34 is substantially poorer, but still achieves some small degree of separation. Finally, a publicly available pretrained contrastive model, denoted as SimCLR-D performs better than ImageNet-ResNet34 but poorer than all of the DIME variants, although it was trained on almost 400,000 histopathology patches with the vast majority being sampled from 35 WSI datasets including 21 TCGA projects [16]; we expect that this inferior performance may be explained by noting that this simCLR model was trained from scratch using a ResNet18 without ImageNet pretraining and it was trained only at the highest resolution of every slide, around 0.25MPP or around 0.5MPP depending on the dataset.

**Table 1:** Comparison of MAP@R on disjoint TCGA test sub-set with inpainting and contrastive approaches.

| Model | MAP@R with TCGA |  |
| --- | --- | --- |
|  | average of the average | weighted average |
| DIME-Inpainting | 0.79 | 0.93 |
| DIME-Inpainting-D | 0.74 | 0.89 |
| DIME-Inpainting-DCT | **0.81** | **0.94** |
| DIME-SimCLR-DC | 0.68 | 0.84 |
| DIME-SimCLR-DCA | 0.59 | 0.75 |
| ImageNet-ResNet34 | 0.44 | 0.66 |
| SimCLR-D [16] | 0.53 | 0.67 |

It is interesting to note that DIME-Inpainting-DCT slightly outperformed DIME-Inpainting in the clustering evaluation, although it is trained with only 3% of WSIs (243,083 patches) used by DIME-Inpainting. DIME-Inpainting-D performs poorer than DIME-Inpainting-DCT despite being trained on the same TCGA diverse dataset, yet is identical to the DIME-Inpainting model in terms of target selection and colour augmentation. Refer to Section 4.1.4 for a description of the differences in model variants. The result comparison suggests that the key factors contributing to the clustering performance improvement with DIME-Inpainting-DCT arise from applying stronger colour augmentation to the input image and choosing the image prior to colour augmentation as the target image.

## 6 VALIDATION EXPERIMENTS, DOWNSTREAM CLASSIFIERS AND APPLICATIONS

This section details additional validation experiments wherein the models trained on TCGA data were applied to H&E-stained WSIs from additional datasets, specifically the publicly available Camelyon16 challenge ^3^, and the AstraZeneca-sponsored Phase III MYS-TIC clinical trial [26].

### 6.1 Tumour Patch Classification on Camelyon16

The downstream application focused on in this section evaluates the ability of the DIME models to capture histology features that are useful for identifying tumour patches. We applied the pretrained models, including ImageNet-ResNet34 and DIME variants as previously described in Section 5.1, to the Camelyon16 dataset that had previously been exhaustively annotated for areas of tumour vs non-tumour. We show the clustering performance on Camelyon16 patch embeddings generated by these pretrained models, and further use them to train a downstream classifier to predict tumorous and non-tumorous patches.

A logistic regression classifier was trained using the 512-dimensional embeddings inferenced from each of the pretrained models. The full-length embedding of each patch is treated as a feature vector, and the pathologists’ annotation of a tumour or non-tumour patch is used for the ground-truth label. In this way the problem is posed as a binary patch-level classification problem. We trained a classifier using the sklearn LogisticRegression tool [27], trained for a maximum of 200 iterations with default parameters.

#### 6.1.1 Camelyon16 dataset

Camelyon16 has been used to identify top-performing computational image analysis systems for the task of automatically detecting metastatic breast cancer in digital WSIs of sentinel lymph nodes. The dataset consists of a total of 400 annotated H&E-stained WSIs split into 270 for training and 130 for testing, and these WSIs were scanned at 40x or approximately 0.25 MPP. Samples in both splits come from two institutions (Radboud University Medical Center and University Medical Center Utrecht) [28].

All WSIs that contain tumour tissues were annotated in a pixel-wise manner by (usually multiple) trained pathologists, while the non-tumour (normal) WSIs remained unannotated. The original Camelyon16 data split is kept for the patch classification. We removed 20 slides from the training dataset and 2 slides from the test dataset because they were not fully annotated. Every slide was patched at four available MPP levels that are closest to 0.5, 1.0, 2.0 and 4.0, resulting in 2,055,957 annotated patches in total, among which 1,353,990 patches were used for training (1,302,439 negatives and 51,551 positives) and 701,967 were used for testing (641,753 negatives and 60,214 positives). Positives refer to tumour patches while negatives refer to non-tumour patches. Such a class definition is used consistently for patch classification problems in this paper.

#### 6.1.2 Clustering results on the Camelyon16 dataset

For displaying UMAP and computing MAP@R metrics on the Camelyon16 dataset, we limited our use to only tumour slides from the test dataset. This was due to CUDA memory limitations making it impractical to compute MAP@R for all patches from the entire test dataset. The quantity of non-tumour and tumour patches in these test tumour slides were relatively well balanced (unlike the entire Camelyon16 test dataset which is heavily biased towards non-tumour patches). The MAP@R metrics are reported in table 2, which come from the average of MAP@R calculated on all patches (both non-tumour and tumour ones), all non-tumour patches, and all tumour patches respectively. The number R is 1023 which is the maximum number of queries supported by the k-nearest-neighbor search algorithm implemented in faiss ^4^. Overall, the two DIME-Inpainting models reported better MAP@R than the two DIME-SimCLR models, particularly for the tumour class, and DIME-Inpainting-DCT slightly outperformed the DIME-Inpainting model despite being trained on only 3% of TCGA slides. This is consistent with the finding in the clustering performances for TCGA test slides in Section 5.2.2. The publicly available pretrained SimCLR-D model reports similar MAP@R values as DIME-SimCLR-DCA, but it is worth noting that the model has seen some Camelyon16 slides during training [16].

**Table 2:** Comparison of MAP@R on tumour slides in Camelyon16 test subset with inpainting and contrastive approaches.

| Model | MAP@R on Camelyon16 test tumour slides |  |  |
| --- | --- | --- | --- |
|  | all patches | non-tumour | tumour |
| ImageNet-ResNet34 | 0.828 | 0.882 | 0.616 |
| DIME-Inpainting | 0.872 | **0.908** | 0.730 |
| DIME-Inpainting-DCT | **0.877** | 0.907 | **0.761** |
| DIME-SimCLR-DC | 0.857 | 0.898 | 0.699 |
| DIME-SimCLR-DCA | 0.859 | 0.905 | 0.684 |
| SimCLR-D [16] | 0.858 | 0.903 | 0.688 |

A UMAP embedding exploration shows that the clustering of patches may be influenced by different scanning conditions and staining intensities. Figure 3 uses different colours to show the distribution of MPP levels of Camelyon16 patches. In the DIME-Inpainting-DCT case, as shown in Figure 3a, the patches from three lowest MPP levels (0.45, 0.46 and 0.49) that are very close in zoom level, however, are assigned to separate clusters of colour black (on the right), dark blue (on the left) and purple (at the bottom) respectively. Some example patches from each of the clusters, for instance, the ones labelled as A, D, F and I in Figure 3a, show that there are distinct variations in staining intensity across the three lowest MPP levels. This suggests that the inpainting approach is very sensitive to colour or staining variations.

**Figure 3:**
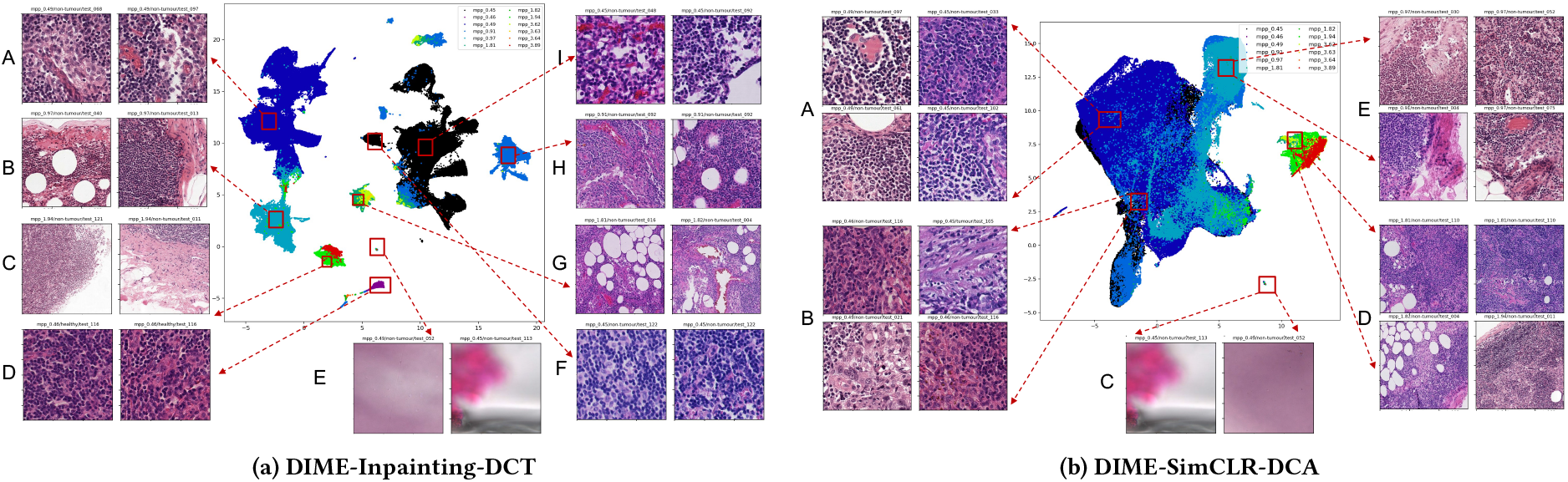
Example Camelyon patches on UMAP coloured by MPP level. The embeddings from DIME-Inpainting-DCT are more sensitive to staining variations than DIME-SimCLR-DCA.

The DIME-SimCLR-DC embeddings generated using the same colour augmentations, however, demonstrated improved robustness to staining variations. Using more aggressive colour augmentation (i.e. the DIME-SimCLR-DCA model variant) the embeddings tended to cluster on similar MPP levels, mostly immune to the influence of staining, as shown in Figure 3b. Two distinct clusters are visible, a larger one mostly comprised of MPP levels ranging from 0.46 to 0.97 and a smaller one covering MPP levels ranging from 1.81 to 3.89. Every group of example patches, labelled as A, B, D and E, shows a good mixture of two or three MPP levels that are very close but have distinctly stained patches. This comparison shows that with appropriate colour augmentations, the SimCLR model variants can be extremely robust to staining variations.

#### 6.1.3 Classification results on the Camelyon16 dataset

Figure 4 shows the Receiver Operating Characteristic (ROC) curves and Precision-Recall (PR) curves of all patch-level classification results on the Camelyon16 dataset. The ImageNet-ResNet34 embeddings are the worst in terms of both area under curve (AUC) and average precision (AP), since the model used was a pretrained ImageNet model that had not been fine-tuned on digital pathology data. The

**Figure 4:**
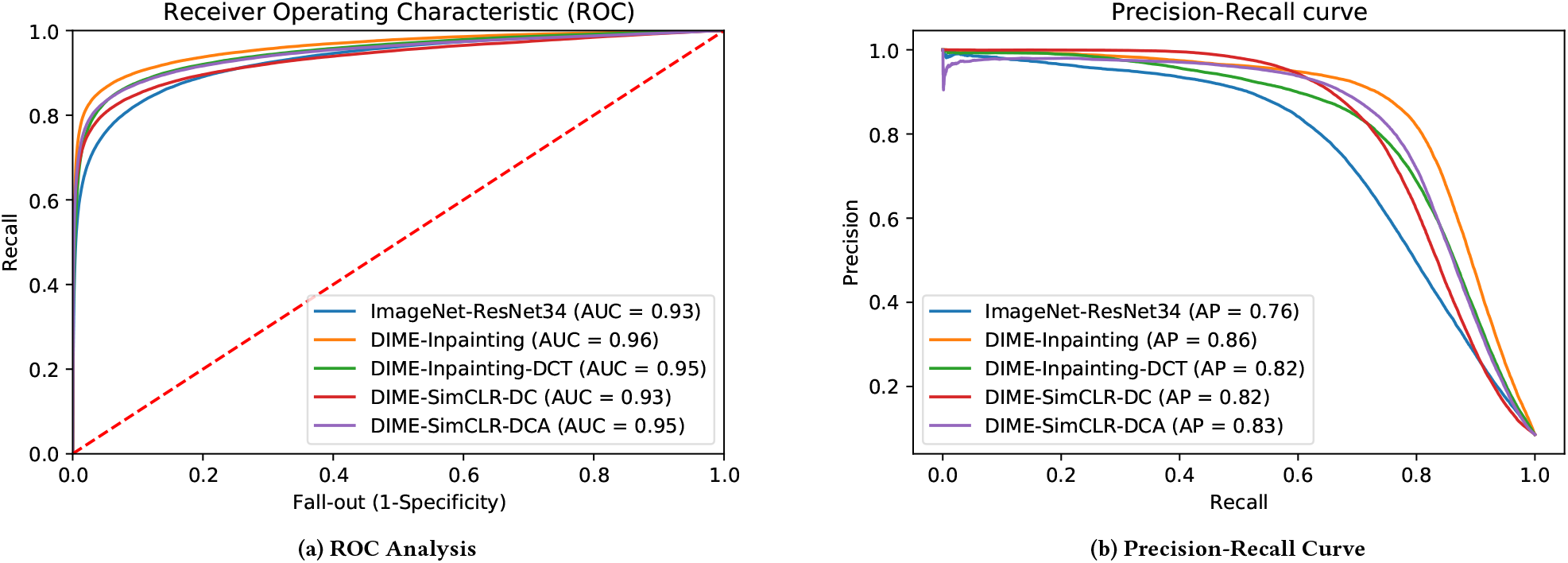
Patch-level classification performance on Camelyon16 test dataset, using embeddings inferenced from ImageNet-ResNet34, DIME-Inpainting, DIME-Inpainting-DCT, DIME-SimCLR-DC, DIME-SimCLR-DCA, respectively.

DIME-Inpainting embeddings show better performance than DIME-Inpainting-DCT and DIME-SimCLR-DC overall. However, this difference in performance is relatively small when considering that the latter two models were trained on 3% of the training data used for the DIME-Inpainting model.

It’s noted that both the Camelyon16 train and test subsets have 12 MPP levels shown in Figure 3 and each subset has images coming from the two institutions (Radboud University Medical Center and University Medical Center Utrecht), meaning that the distribution of staining variations in the train subset should be very similar to that in the test subset. This helps the classifier to generalise well to the test subset even though the embeddings from inpainting models are sensitive to staining variations.

Given these trained linear patch classifiers, a further step is taken to predict the slide-level label (i.e., tumour or non-tumour), by preforming inference over patches for every testing slide, generating a tumour probability heatmap and reporting the maximum value in the heatmap as the slide-level tumour prediction. We compare with the three results reported by [13] as shown in Table 3. Max-pooling and DSMIL are multiple-instance learning (MIL) models that uses patch embeddings to predict the label of the testing slides. The patch embeddings used in [13] are inferenced from a simCLR model which was trained on Camelyon16 training slides. In our case, however, the DIME-SimCLR model, as the feature extractor, was trained only with TCGA slides. The upper-bound performance, reported in [13] is the fully-supervised model which trains a deep model with labelled patches. Although our patch classifier is simply a linear model without performing any upsampling technique, the WSI classification results using DIME-SimCLR-DCA embeddings of 20x (close to 0.5 MPP) patches achieves very competitive results, matching the accuracy performance of the simCLR-max-pooling method in [13], with a classification AUC gap smaller than 2%. This suggests that the quality of Camelyon16 patch embeddings is competitive even without using Camelyon slides for fine-tuning the feature extractor.

**Table 3:** Comparison of slide-level classification on Came-lyon16 test slides. DIME-SimCLR-DCA is pretrained only on TCGA slides, whereas all other methods were pretrained on Camelyon16 slides.

| Features | Slide-level Camelyon16 classification |  |
| --- | --- | --- |
|  | Accuracy | AUC |
| DIME-SimCLR-DCA | 0.8295 | 0.8431 |
| Max-pooling [13] | 0.8295 | 0.8641 |
| DSMIL [13] | 0.8682 | 0.8944 |
| Fully-supervised [13] | **0.9147** | **0.9362** |

### 6.2 Tumour patch classification on the MYSTIC Phase III clinical trial dataset

The downstream application focused on in this section is the classification of tumour vs non-tumour patches of H&E-stained WSIs from a clinical trial dataset using DIME embeddings. We applied the DIME pipeline, trained only on TCGA diagnostic slides as previously described, to H&E-stained WSIs from MYSTIC (NCT02453282), an AstraZeneca-sponsored Phase III randomized clinical trial of durvalumab or durvalumab plus tremelimumab vs chemotherapy in first line treatment of metastatic non-small cell lung cancer (NSCLC) [26]. A total of 534 H&E-stained WSIs from biopsies obtained prior to the start of therapy were used for this analysis. Areas of tumour were annotated as polygons by 2 trained pathologists, in a similar manner as was done for the Camelyon16 dataset.

To investigate the effect of colour normalisation on clustering and tumour patch classification, the colour normalisation procedure described in [29] was applied to the MYSTIC slides.

#### 6.2.1 Embedding exploration for MYSTIC

Figure 5 shows a comparison of the UMAPs prior to and after color normalisation on 20 MYSTIC WSIs that were randomly selected from the dataset. As we expected, the UMAPs after colour normalisation display more overlap among different patients in comparison to the ones prior to colour normalisation. The difference between pre- and post-colour normalisation is particularly obvious in the cases of DIME-Inpatinting and DIME-Inpainting-DCT model variants. This further proves that some embedding clusters created by the in-painting approach prior to colour normalisation are formed due to the variations in staining. On the other hand, we see that the two UMAPs from DIME-SimCLR-DCA are very similar to each other in terms of the degree of overlap among patients, indicating that the contrastive approach can be robust to staining variation given that there has been aggressive colour augmentations applied to the training images.

**Figure 5:**
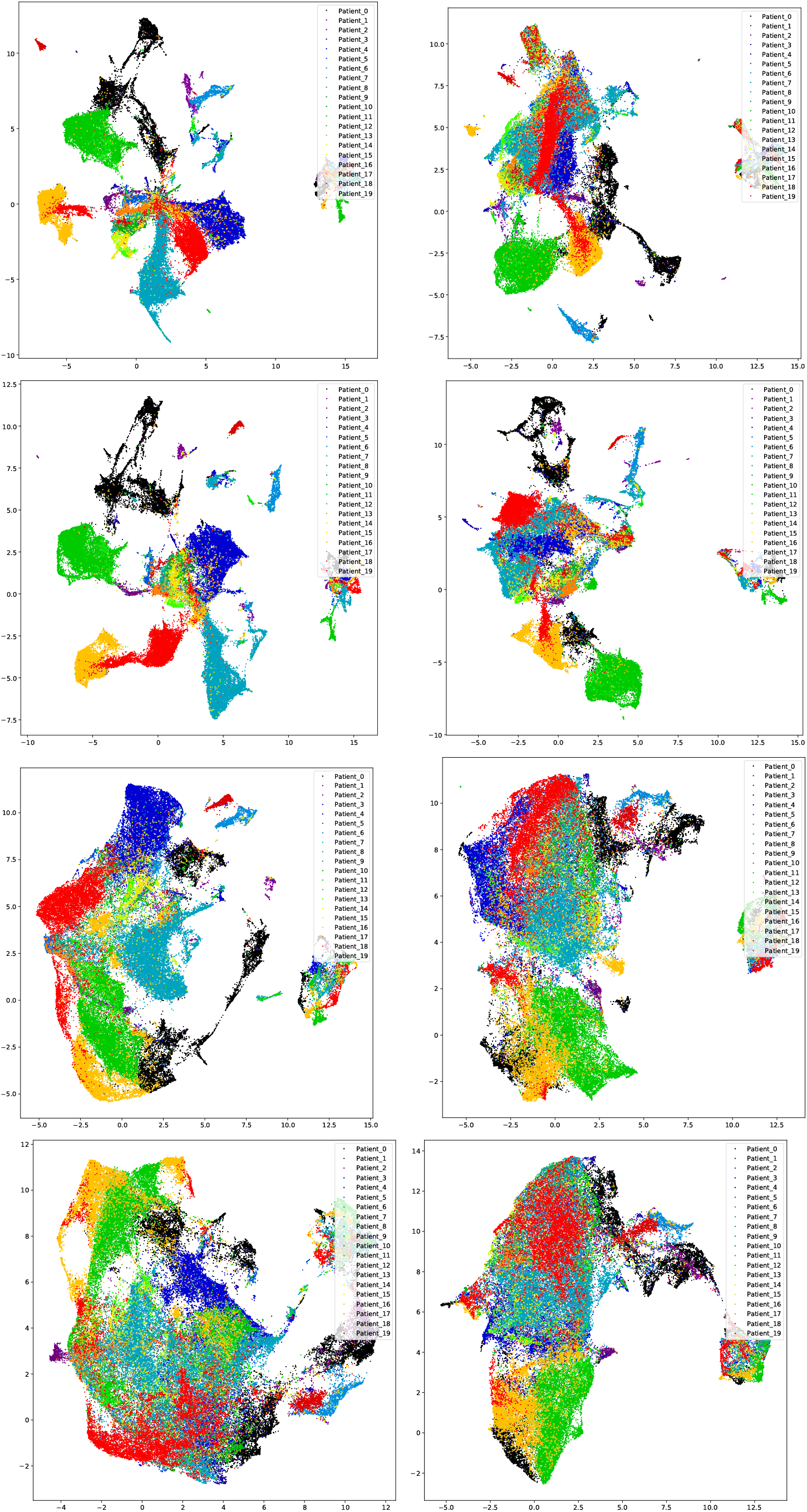
A comparison of UMAPs for embeddings generated with the DIME-Inpainting (top row), DIME-Inpainting-DCT (second row), DIME-SimCLR-DC (third row) and DIME-SimCLR-DCT model variants, for original (left) and colour-normalised (right) MYSTIC image patches.

A deeper dive into the UMAP from DIME-Inpainting, as shown in Figure 6, helps understand more why some patients’ patch embeddings are merged into one large cluster while others remain separated. The green, orange and the adjacent black islands are well separately from the largest cluster even after colour normalisation. This is because the three separated islands comprise all tumour patches, as shown in Figure 6D, E and H, while the majority of the largest cluster consists of non-tumour patches, according to the pathologist annotations. By contrast, some tumour patches from Patient 7, like the two in Figure 6B, are close to the non-tumour ones (such as Figure 6C) in the largest cluster at the UMAP space, suggesting that these tumour patches are more difficult to tell from the non-tumour ones in terms of their morphological features. The three sets of example patches from Patient 0 and Patient 6, shown in Figure 6I, G and F, demonstrate that non-tumour tissues of different morphological characteristics can be clustered separately.

**Figure 6:**
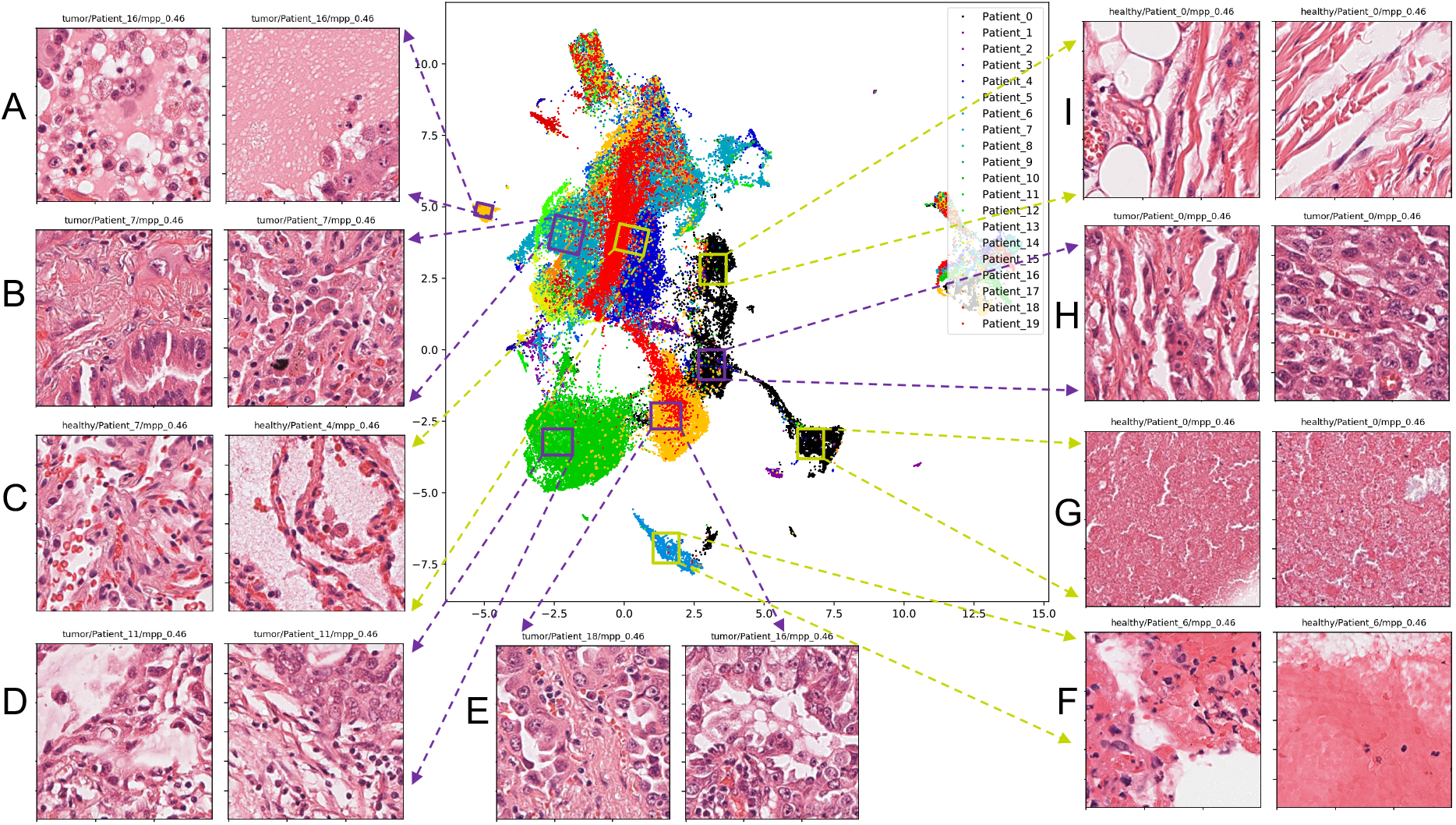
UMAP for non-tumour/tumour exploration of embeddings generated with the DIME-Inpainting and colournormalised MYSTIC patches.

#### 6.2.2 Classification results on the MYSTIC dataset

The 534 annotated slides from the MYSTIC dataset were split at the slide level into training (60%) and testing (40%) subsets, and every slide was divided into patches at four available MPP levels (0.46, 0.92, 1.84 and 3.68). This resulted in 595,178 annotated patches in total, among which 595,178 patches were for training (306,391 negatives and 288,787 positives) and 324,533 were for testing (166,289 negatives and 158,244 positives). A LR classifier as described in 6.1 was trained using the embeddings from these annotated patches generated by every pretrained model of interest.

Figure 7 shows ROC and Precision-Recall curves on the MYSTIC testing subset slides prior to colour normalisation. The embeddings produced by the ImageNet-ResNet34 show the worst overall performance on ROC AUC and AP. The DIME-Inpainting embeddings lead to the best performing classification, with the DIME-Inpainting-DCT and DIME-SimCLR-DCA embeddings coming second and the DIME-SimCLR-DC model close behind. However, the DIME-Inpainting-DCT and the DIME-SimCLR-DC embeddings resulted in only slightly lower classification performance despite being trained on only 3% of the data used to train DIME-Inpainting.

**Figure 7:**
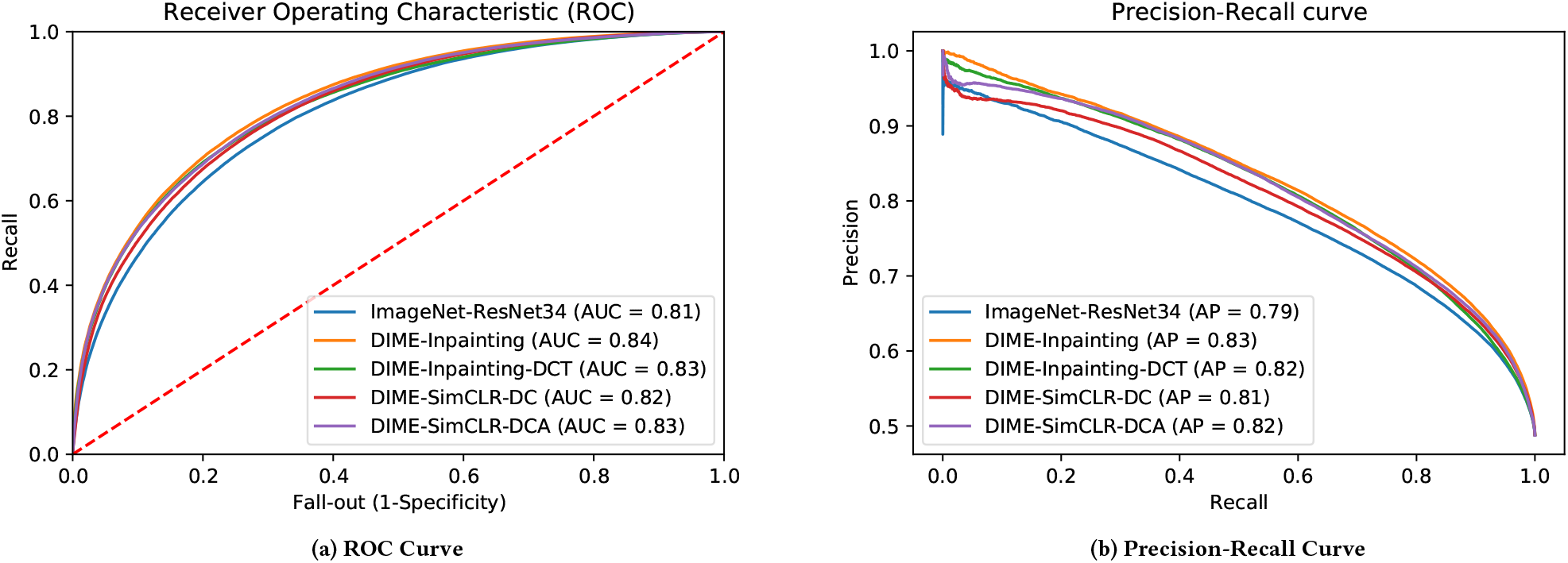
The comparison of ImageNet-ResNet34 and DIME variants for the patch-level classification on MYSTIC dataset.

### 6.3 Automated annotation application

With a working patch classifier, it then becomes possible to implement an automated annotation pipeline. This pipeline is triggered as new imaging data are acquired, and uses DIME to generate patch embeddings before feeding these embeddings in to our pretrained patch classifier. In this way it is possible to produce tumour patch annotated images in a completely automated fashion.

Figure 8 shows an example of this in action, with side-by-side annotated images; one by a pathologist, and one using our patch classifier with a cutoff threshold of 0.5. As a practical matter we improve the output by utilising relevant spatial information; e.g. by smoothing predictions across adjacent patches with a median filter. Here we show a demonstration on Mystic, but this automated annotation procedure can be applied to any such dataset (large or small).

**Figure 8:**
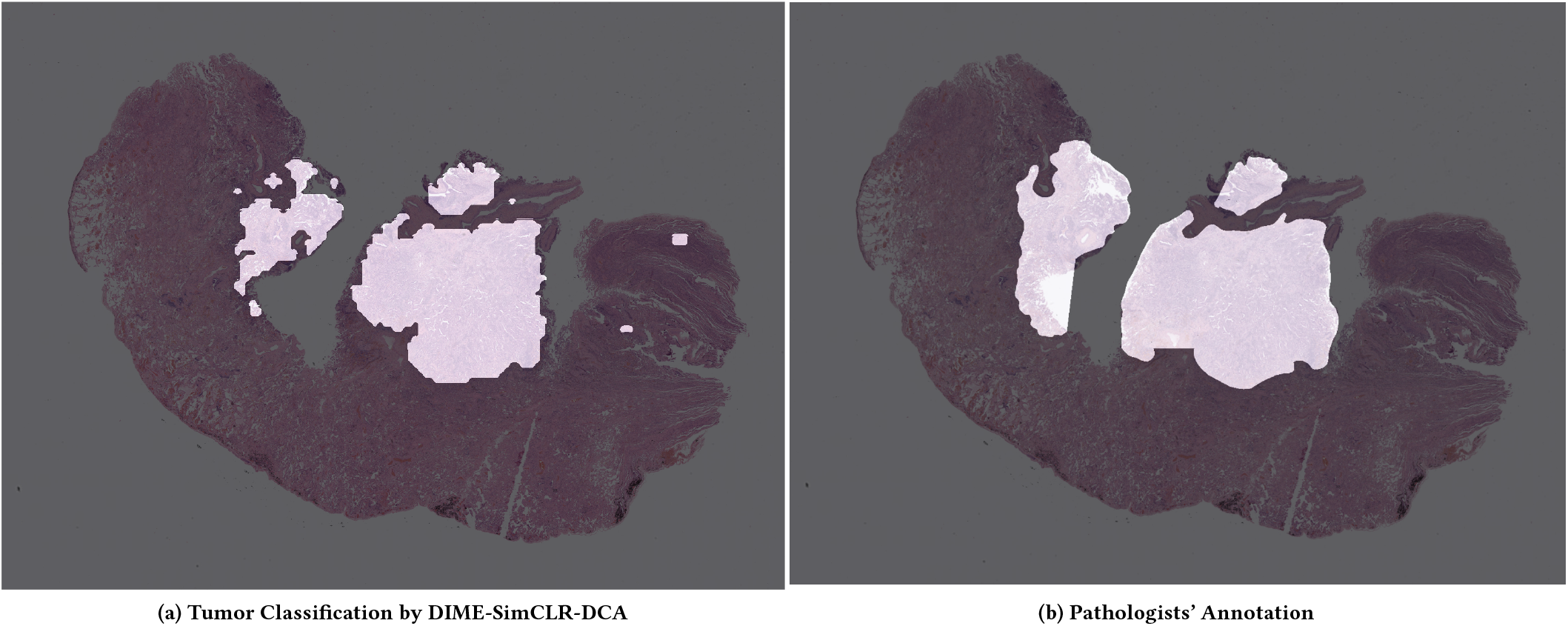
The comparison of tumour classification and annotation for a slide from the MYSTIC dataset.

An effective automated annotation application is increasingly valuable for drug development as ever larger imaging datasets become available. Expert pathologists are a scarce resource and their time is not best spent hand labelling hundreds or thousands of images that can be largely done by an automated application. In practice it is useful to have an expert validate small quantities of the automatically annotated images, and we believe that future improvements to the automated annotation pipeline would involve using confidence estimates to draw attention to the least certain regions, and potentially also human-in-the-loop training regimes to bootstrap performance while most efficiently making use of expert pathologists’ time.

### 6.4 Quality control application

One proposed application of DIME embeddings is quality control, whereby poor quality images or image patches are automatically identified and removed from further processing. For this be successful the embeddings should cluster on particular image artefacts or corruptions, regardless of the particular tissue or disease type. To explore this possibility, we examine clusters of embeddings produced by the DIME-Inpainting model for TCGA with reference to the patches from which these embeddings were produced.

Figure 9 shows the two dimensional downprojection of the 512 dimensional embeddings produced for the TCGA training dataset via UMAP. In addition we show example patches randomly selected from within the regions where multiple tissues or disease types cluster together.

While the down-projected embeddings show good clustering according to cancer type and MPP level, there are some noteworthy cases of overlapping embeddings. We explored the patches corresponding to these embeddings found in overlapping regions by visual inspection of a few selected cases.

**Figure 9:**
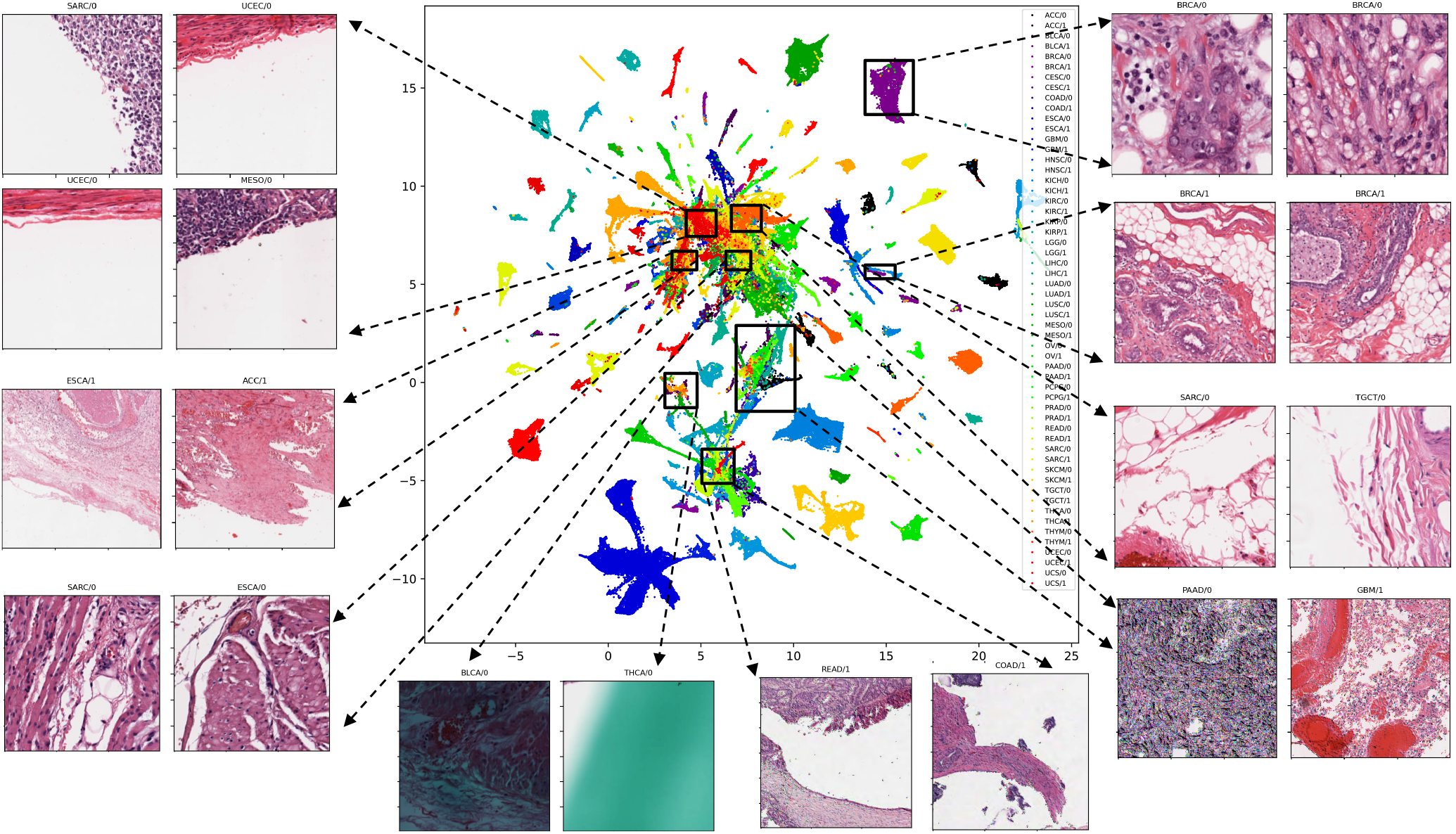
UMAP visualization of TCGA test images from DIME-Inpainting.

As Figure 9 shows, the largest overlapping clusters are dominated by edge patches that have a large proportion of background. It would therefore be trivial to exclude such cases from analysis by setting a higher background threshold cutoff in the early stage of the DIME pipeline. However, filtering in this way would reduce the quantity of edge tissue analysed (which pathologists have expressed an explicit desire to focus on in some cases), so there is a natural trade-off between background cutoff threshold and patch size that could be optimised for particular applications.

Some other areas with overlap reveal common issues with image quality, such as image corruption, blurring, or pen marking. The overlapping regions serve to illustrate two points: first, that the MAP@R objective metrics based on these labels are necessarily imperfect, and second, that the overlapping regions are usually signs of good behaviour on the part of the model, in that genuine patch similarities have been identified.

The clustering across tissue types on common image corruptions gives a strong indication that the DIME embeddings can be used for a quality control application. A quality control application could be built on top of the DIME embeddings in one of several ways. For example, most simply the regions of the embedding space corresponding to particular artefacts could be used to label any new patch embeddings with the corresponding artefact. Alternatively more complex implementations could utilise clustered regions to provide pseudo-labels for a new model to be trained before it is applied to additional datasets.

## 7 DISCUSSION

DIME has many potential applications across the spectrum of drug development. By serving as a searchable WSI repository and enabling the classification of H&E-stained images, DIME can be used for automated annotations, image quality control, and exploratory analyses. It is a useful tool for pathologists, researchers, and clinicians to streamline their workflows and to gain novel insights into clinical trial imaging datasets, with demonstrated performance on real world global clinical samples used in drug development.

DIME is able to classify similar H&E images, enabling object identification and thereby serving as a proxy for automated annotation of features including tumour epithelium, tumour-associated stroma, and varying amounts of inflammation. It can provide a second opinion to pathologists who, when faced with a challenging case, can draw on information from similar images to help inform diagnosis or feature assessment. One particularly useful practical application is artefact identification. By identifying patches that contain artefacts such as pen marks or folds, as highlighted in Figure 9, DIME can identify images that may require special treatment in pathology workflows, such as rescanning, restaining, or patch exclusion prior to use in additional applications which may be sensitive to artifacts.

By classifying images within and between datasets, DIME can be used for biological hypothesis generation. For example, DIME could be used to identify morphologic features contained in H&E images which are associated with genomic alterations including gene or pathway aberrations present in a subset of patients. These features may be too subtle to be readily distinguishable by the eye but could be picked up by AI. Classification of mutations by DIME could replace more expensive (and time-consuming) genomic assays and may be useful to match patients to the specific treatment most likely to work for them.

Classification by DIME could also be used in exploratory analyses to identify features associated with drug response or resistance - assessed from pre-treatment biopsy samples or by looking at changes that occur in a patient’s tumour by the comparison of both pre- and on-treatment biopsy samples. These observations could be used for therapy selection or for patient monitoring while on-treatment. Manual review of DIME classifications by pathologists may also enable the identification of TME features associated with response or resistance categories which can be explored further using other imaging approaches or genomic or other assays.

Additionally, while it is widely accepted that cancer is not a single disease and the identification of cancer subtypes is important for defining prognosis and treatment, cancer classification systems are still largely based on the tissue of origin. DIME provides an opportunity to classify tumour features across organ locations in a data-driven way to discover pan-cancer pathomorphologic phenotypes. Findings will enable precision medicine approaches to target therapies to subjects that previously may not have been considered for a therapy because of the organ location of their disease.

Finally, while DIME as applied here has been restricted to images from H&E-stained biopsies, the DIME approach has the potential to be applied to additional imaging modalities, including IHC, multiplex technologies (e.g. multiplex IHC, multiplex immunofluorescence, imaging mass cytometry), and radiology images (e.g. CT, PET, MRI). For each new application, DIME has the potential to make significant contributions to the improvement of medical diagnostic workflows and the discovery of novel biology to advance drug development.

## 8 CONCLUSION

In this work we introduced the DIME pipeline which effectively converts WSIs into patch embeddings that may be explored to derive novel insights into underlying biology or utilized to improve pathology or downstream analysis workflows. The embeddings are trained without the need of labels or metadata, and demonstrate useful properties allowing downstream applications to be built such as similar image search, automated annotation, quality control, and training predictive models.

We explored the relative strengths of SimCLR and inpainting for embedding generation, finding inpainting to produce embeddings ultimately better at tissue/disease-type clustering, whereas SimCLR is more robust to colour variations introduced by staining and scanning and hence may not require image colour normalisation at inference time in a practical application.

Lastly, we applied the DIME pipeline to H&E-stained WSIs from a immuno-oncology clinical trial and demonstrated how DIME may be used in a tumour classification automated annotation pipeline, potentially expediting pathologist workflows and enabling a deeper understanding of how and why therapies succeed or fail. The embeddings from DIME extract additional pathomorphologic information from H&E-stained images which may also be used to train predictive models of therapy response. These models could be used to advance drug development by helping clinicians identify which therapy is right for which patient as part of a personalized-medicine approach to cancer treatment.

## ACKNOWLEDGMENTS

Acknowledgments should be inserted at the end of the paper, before the references, not as a footnote to the title. Use the unnumbered Acknowledgements Head style for the Acknowledgments heading.

## SUPPLEMENTARY MATERIAL

### Details of TCGA dataset

Figure 10 shows the distribution of MPPs across the generated TCGA images.

**Figure 10:**
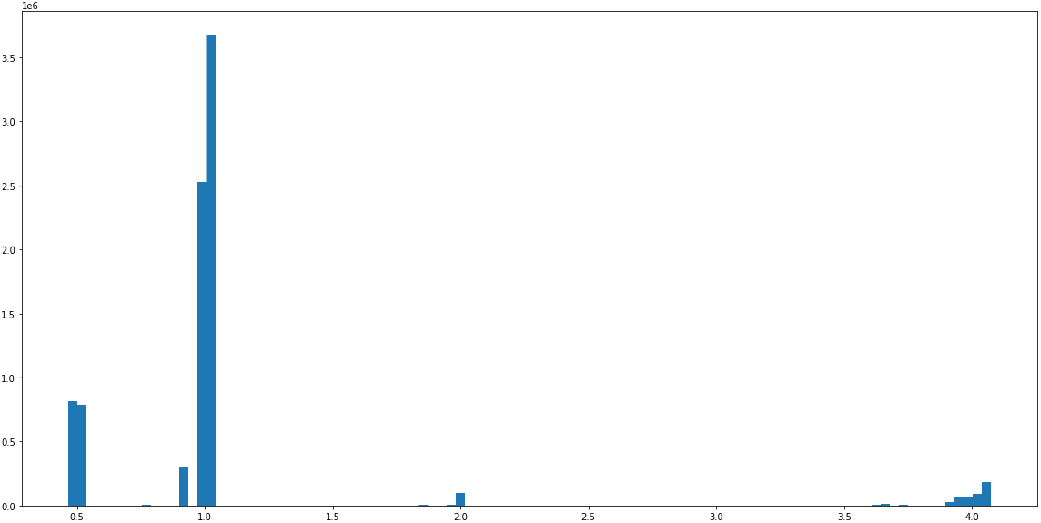
Distribution of MPPs across generated TCGA image patches.

Figure 11 shows the total quantity of patches extracted per TCGA project and per MPP level across train, validation and test splits.

**Figure 11:**
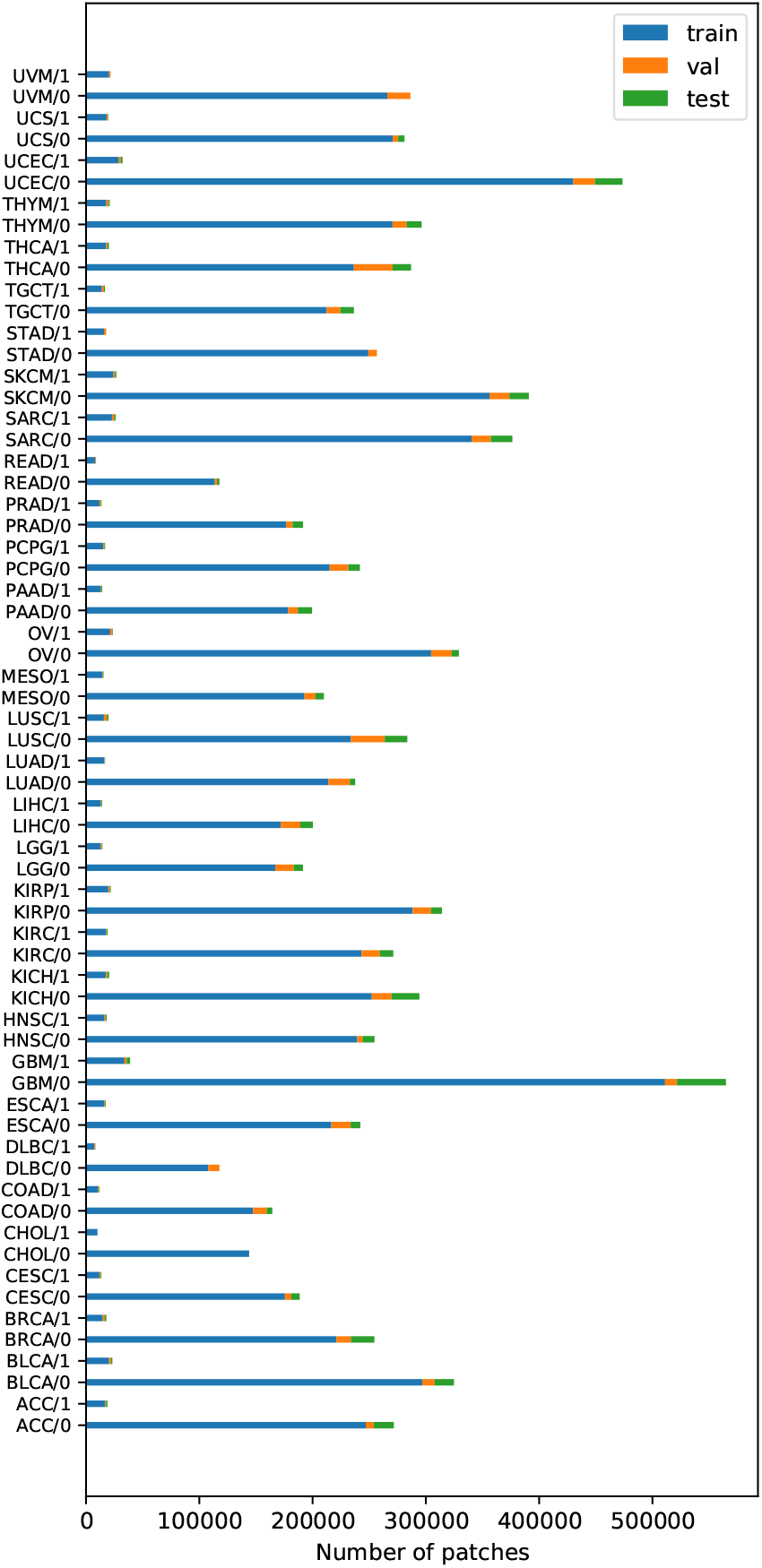
Distribution of patches per TCGA project.

### Details of training setup

#### Inpainting setup

In the inpainting approach, every image patch is filled with a random number (ranging between 3 and 7) of black boxes. Every black box takes a minimum of 10% and a maximum of 30% of the image size. The inpainting models were trained on a 4-GPU machine with a batch size of 32 patches per GPU. The Adam optimizer is used in combination with Leslie Smith’s 1-cycle policy [30] with a learning rate ranging between 1e-6 and 1e-4. We train each network without freezing any layers (fine-tuning).

#### SimCLR setup

To train a SimCLR model, we used the Adam optimizer combined with initial learning rate of 1e-5 and Cosine annealing with initial 10 warmup epochs. For the contrastive loss, we use NT-Xent loss and temperature scaling of 0.5. Nvidia-apex is enabled for mixed precision training to accelerate the training and the batch size is 188 patches per GPU (i.e., the number of unique slide labels * the number of MPP levels).

#### Early Stopping

Figure 12 shows the change in MAP@R on the validation subset of the DIME-Inpainting-DCT and DIME-SimCLR-DC models, which shows that convergence was observable after around 200 rounds. The DIME-Inpainting model was trained for 1600 rounds and, as shown in Figure 13, did not yet fully converge.

**Figure 12:**
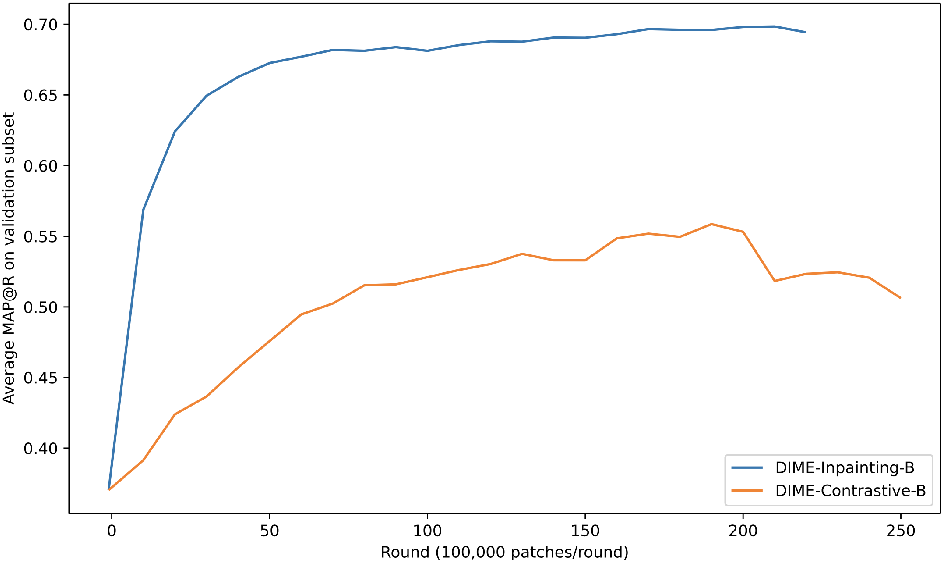
Validation MAP@R from DIME-Inpainting-B and DIME-Contrastive-B models.

**Figure 13:**
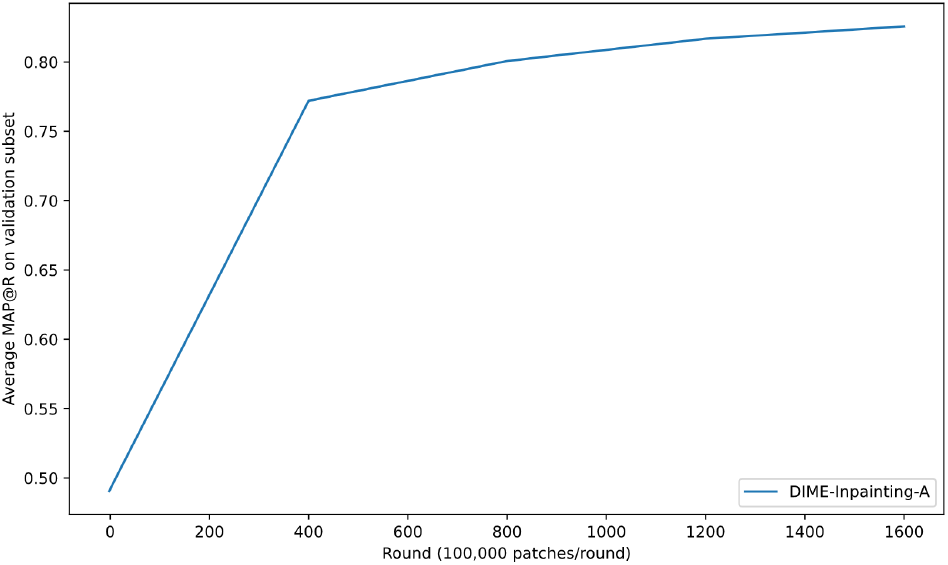
Validation MAP@R from DIME-Inpainting-A model.

### MYSTIC patch classification performance with reduced training data

We explored the effect of training set size by adjusting the ratio of data used for training and testing of MYSTIC patch classification models. The change in AUC and AP with decreasing training set size is shown in Figure 14.

**Figure 14:**
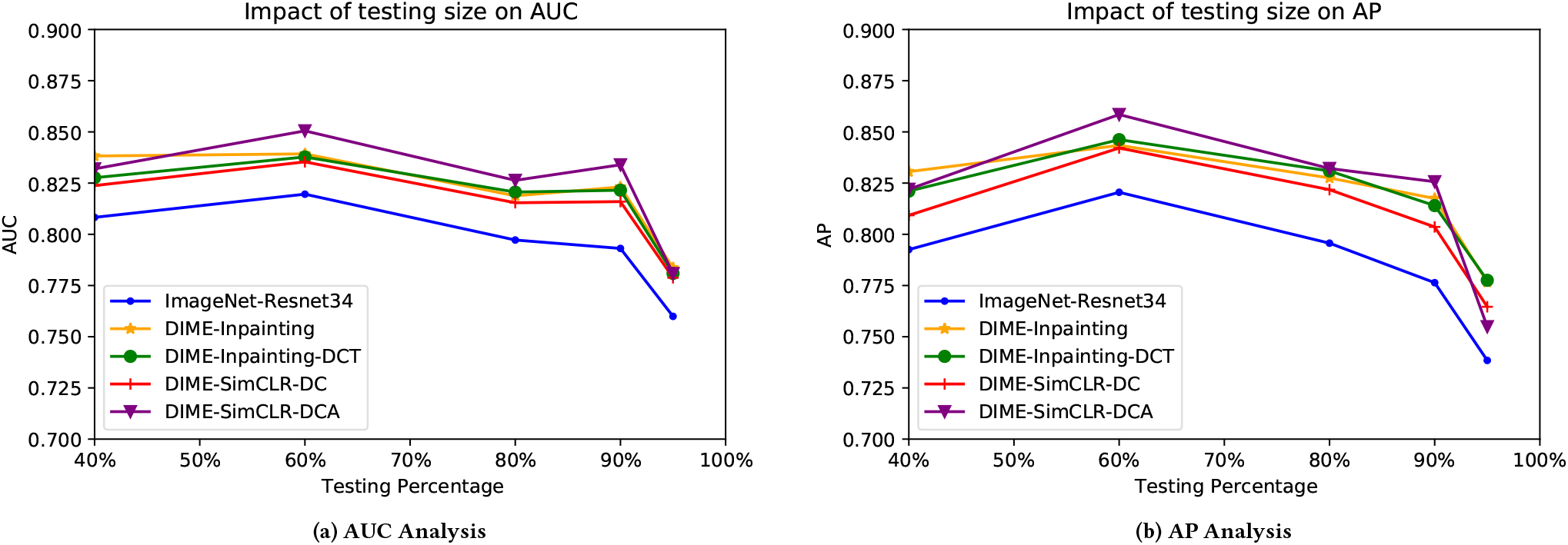
Change in AUC and AP as the MYSTIC testing size increases, reported from ImageNet-ResNet34 and DIME variants.

We note that it is only after we reduced the training set to 5% of total annotated slides (i.e. training on 27 annotated slides) that we observe a noticeable deterioration in patch classification performance. Even with this deterioration, the AUC remains above 0.77 and the AP above 0.75. These are encouraging results indicating that the DIME embeddings are able to capture the key differences between tumorous and non-tumorous patches, and hence only a handful of annotated slides, are required to produce an effective classifier for automated annotation. This implies that similar classifiers could be trained for a variety of tissue and disease-types with very small quantities of initially annotated data (which are expensive to obtain).

## Footnotes

1 https://portal.gdc.cancer.gov/

2 The pretrained simCLR model is available to download from https://github.com/ozanciga/self-supervised-histopathology/releases/tag/tenpercent

3 https://grand-challenge-public.s3.amazonaws.com/f/challenge/80/105788c6-176a-4dc3-89cf-62f4f37d1484/camelyon16_readme.md

4 https://github.com/facebookresearch/faiss

